# Full-Length Structural Modeling of Mitofusins with AlphaFold Reveals a Novel Cross-Type Dimerization and Insights into Oligomerization

**DOI:** 10.64898/2026.04.10.717648

**Authors:** Raphaëlle Versini, Marc Baaden, Alexandre M.J.J. Bonvin, Patrick F.J. Fuchs, Antoine Taly

## Abstract

Mitochondrial dynamics, involving fission and fusion, are critical for the maintenance, function, distribution, and inheritance of mitochondria, allowing their morphology to adapt to the cell’s physiological needs. Mitofusins, large GTPase transmembrane proteins, play a role in this process by driving the tethering and fusion of mitochondrial outer membranes. Dysfunction in mitofusins has been associated with neurodegenerative diseases such as Parkinson’s, Alzheimer’s, Huntington’s, and Charcot-Marie-Tooth type 2A, as well as various cancers, where their dysregulated expression influences cell proliferation, invasion, and chemotherapy resistance. Despite their importance, the precise molecular mechanisms underlying mitofusin-mediated fusion remain unclear and require further structural elucidation. In fact, no complete high-resolution structures exist for mitofusins, including their yeast homolog, Fzo1.

Here, we generated and analyzed full-length structural models of mitofusins using AlphaFold. Monomeric, dimeric, and tetrameric assemblies were produced, including complexes with fusion partners Ugo1 and SLC25A46. While AlphaFold predicted limited conformational diversity for isolated monomers, structural variability emerged for predicted homo- and hetero-oligomers. Notably, our models reveal a previously undescribed cross-type dimerization mode involving interactions between heptad repeat domains, not reported in current experimental structures. Comparison with recently resolved experimental data further supports the structural relevance of this interface. These full-length models allowed us to propose a new hypothetical mechanism of outer mitochondrial membrane fusion.

## 1 Introduction

Mitochondria are unique organelles with two distinct membranes whose most prominent role is to provide the cell with metabolic energy in the form of ATP generated by oxidative phosphorylation. Mitochondria form a dynamic network within cells, undergoing constant fusion and fission events that occur independently of the host cell’s division. These processes are crucial for shaping mitochondrial dynamics and ensuring their maintenance, functionality, distribution, and inheritance [1].

Mitofusins are large GTPase transmembrane proteins involved in the tethering to and fusion of mitochondrial outer membranes (OM) [2]. Among those, Mfn1 and Mfn2 are the two mitofusins found in mammals [3, 4] and Fzo1 (*Fuzzy Onion 1*) is the *Saccharomyces cerevisiae* homolog [5]. Structures of Mfn1 (PDB: 5YEW, 5GOF, 5GOM), Mfn2 (PDB: 6JFK) and Fzo1 (PDB: 9KFE, 9KFF, 9KFD) were partially solved, but without the transmembrane domain and most of the heptad repeat (HR) domains [6, 7]. Mitochondrial inner membrane fusion and cristea organization are mediated by human OPA1 (*Optic Atrophy 1*) [8] and yeast Mgm1 (*Mitochondrial Genome Maintenance 1*) [9].

Mitochondrial fusion dysfunction can cause neurodegenerative diseases such as Parkinson, Alzheimer, and Huntington [10, 11] as well as cancer [12, 13]. Several studies have linked Mfn1 and Mfn2 to various cancers, including breast [14], liver [15], lung [16], cervical [17], and colon [18] cancers. Their dysregulated expression was associated with increased cell proliferation, invasion, and resistance to chemotherapy. Regulation of mitofusin activity has been shown in preclinical studies to reduce cancer cell growth and spread. Furthermore, studies have demonstrated that mutations in Mfn2 drive the onset and progression of muscular dystrophies, such as Charcot-Marie-Tooth type 2A, the most prevalent form of axonal CMT disease [19, 20]. The exact mechanisms through which mitofusins contribute to cancer and neurodegenerative diseases, as well as the molecular details of the fusion process they mediate, remain to be elucidated through further structural studies.

Ugo1 is a modified mitochondrial solute carrier embedded in the outer mitochondrial membrane shown to be required for outer mitochondrial membrane fusion. Its specific role may not yet be fully understood. Indeed, Sesaki et al., 2001 [21] demonstrated that mutants lacking Ugo1 exhibit numerous small mitochondrial fragments, contrasting with the few long tubular-shaped mitochondria observed in wild-type cells. Furthermore, it was outlined that residues 630–703 of the N-terminal cytoplasmic domain and residues 756–843 of the C-terminal domain were binding to Fzo1 [22]. The human orthologue to Ugo1 would be the member of the mitochondrial solute carrier family SLC25A46 [23, 24]. SLC25A46 has been shown to participate in both upregulation [25] and downregulation [26, 23] of mitochondrial fusion. Hence, the role of the protein is argued to be associated with an overall regulation of mitochondrial dynamics. So far, Ugo1 has only been shown to upregulate mitochondrial fusion [21, 27]. Neither of Ugo1 nor SLC25A46 have an identified substrate, despite their apparent belonging to the solute carrier family [25, 24].

The structure of Mfn1 has been partially elucidated through X-ray diffraction, revealing its monomeric form in 2017 (see Figure 1A and C, [6]. Additionally, two dimeric conformations were identified in 2017 (see Figure 1D) [6] and 2018 (see Figure 1E) [7]. The overall monomeric structure of the protein was consistent with characteristics of the dynamin superfamily, and more specifically the BDLP protein [6, 30].

**Figure 1:**
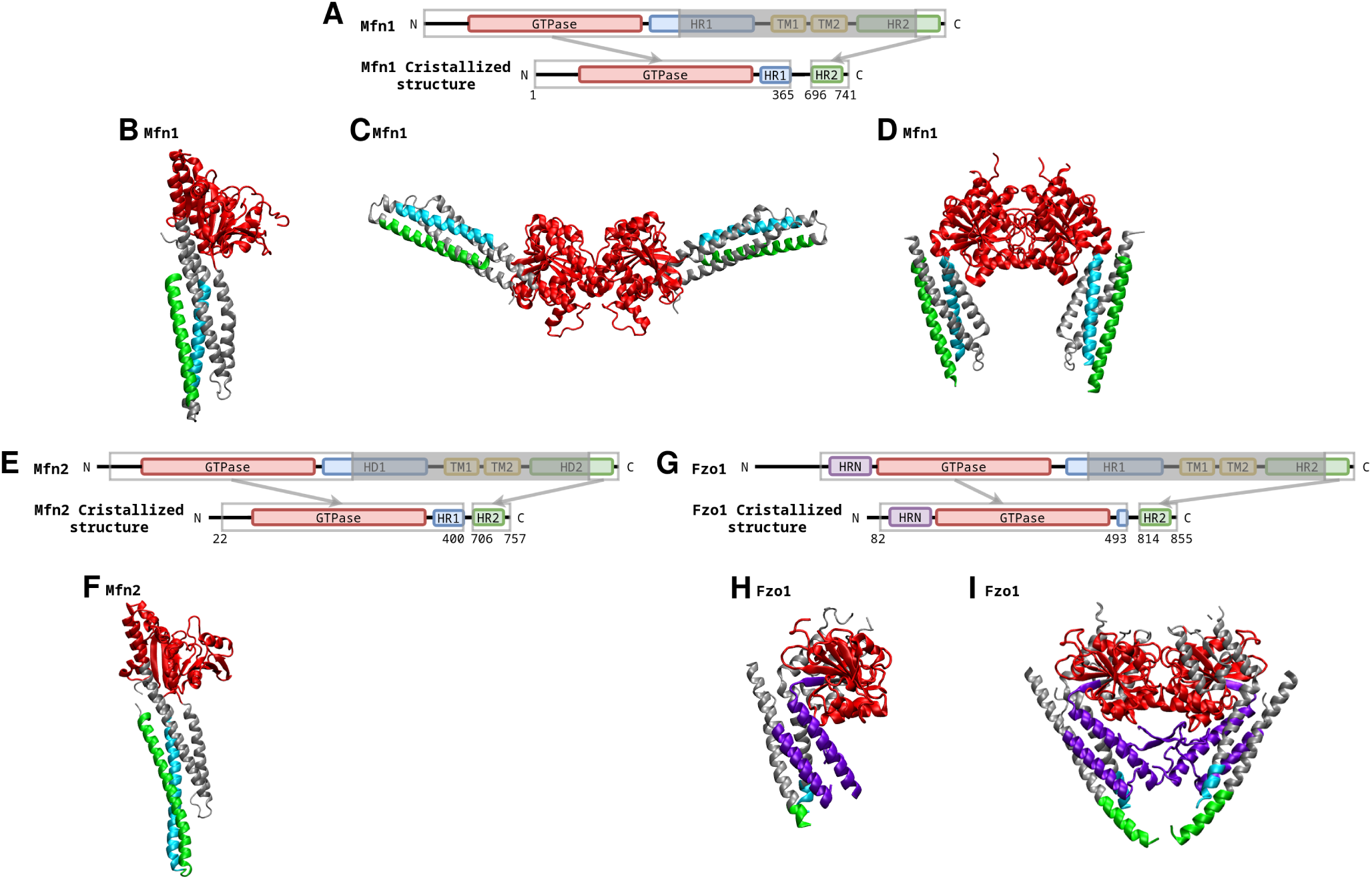
Solved Mitofusins structures. (A) Schematic representation showing the organization of Mfn1 based on full-length Mfn1. GTPase, GTPase domain in red; HR1/2, heptad repeat 1/2 in blue/green; TM, transmembrane region in orange. Borders of each element are indicated by residue numbers. (B) Partial monomer of Mfn1 (PDB: 5GOF) [6]. (C) Dimer conformation of Mfn1 (PDB: 5GOM) [6]. (D) Closed dimer conformation of Mfn1 (PDB: 5YEW) [7]. For (B), (C) and (D) the domains are colored according to (A). In grey is represented the portions of the sequences not assigned to a specific domain. (E) Schematic representation showing the organization of MFN2 based on full-length MFN2. G domain, GTPase domain; HD1/2, helical domain 1/2; TM, transmembrane region. Borders of each element are indicated by residue numbers. (F) Structure of the partial monomer of Mfn2 (PDB: 6JFK) [28], the domains are colored according to (E). In grey is represented the portions of the sequences not assigned to a specific domain. (G) Schematic representation showing the organization of Fzo1 based on full-length Fzo1. GTPase, GTPase domain in red; HR1/2, heptad repeat 1/2 in blue/green; TM, transmembrane region in orange. Borders of each element are indicated by residue numbers. (H) Partial monomer of Fzo1 (PDB: 9KFE) [29]. (I) Dimer conformation of Fzo1 (PDB: 9KFD) [29]. For both (I) and (J), the domains are colored according to (G). In grey is represented the portions of the sequences not assigned to a specific domain.

The structures highlighted the significance of G domain dimerization, which is controlled by guanine nucleotides, in the process of membrane fusion that is facilitated by mitofusin. Upon GTP loading, there is a possibility of inducing a conformational change, transitioning from the ‘closed’ state, characterized by tethering constraints, to the ‘open’ state, which permits tethering [6]. A second, dimeric structure, showed a rearrangement of its internal structure, specifically between the helix bundle and the GTPase domain [7]. In 2019, the structure of Mfn2 was partially solved, as well, in a monomeric conformation (see Figure 1B and F) [28]. The partial structure is very similar to the partial structure of Mfn1 and of the BDLP GTPase domain and helix bundle. Like other dynamin superfamily members, MFN2 possesses key characteristics such as domain organization and G domain-mediated dimerization.

Recently, Huang et al. reported a new structure of yeast Fzo1 (see Figure 1G-I) based on cryoelectron microscopy and biochemical analyses [29]. This structure solves mainly the HRN and GTPase domains. Importantly, this work proposes a revised fusion mechanism in which GTP loading promotes trans dimer formation, followed by a conformational transition upon GTP hydrolysis that brings opposing membranes into close proximity. These studies reinforced structural insights into the mitofusin mechanism of action. They also highlighted the remaining gaps, particularly concerning the membrane-embedded regions, that are likely critical for understanding the overall membrane fusion mechanism.

The possible conformations of mitofusins have been studied using a combination of techniques, ranging from homology modeling to threading [31]. Given the similarity between the mitofusins and BDLP, a few models were built based on the solved structures of BDLP [30, 32], and it was proposed that mitofusins change conformation from an open to a closed state. In 2013 [33], Anton and collaborator constructed an initial model of the yeast mitofusin Fzo1 by using the closed conformation of BDLP as a template, subsequently followed by another model produced by De Vecchis in 2017, which improved the overall structural definition, as well as the transmembrane domain conformation [34]. Importantly, this revised model is consistent with experimental data and MD simulations in explicit membranes [34]. The open conformation was built in 2019 from the closed conformation [35]. Within this framework, the overall structure of Fzo1 is modeled, offering insights into its helical segments. Notably, HR1 and HR2 are not continuous helices, but exhibit bends, similar to flexible hinges observed in BDLP [30, 32]. Models of Mfn2 were proposed in 2016 [36] and 2017 [37] using similar methodologies. These models suggested that a conformational change, involving the unfolding of the HR2 domain, occurs prior to tethering and fusion.

Recently, we worked on a comprehensive study focused on the TM domain of Fzo1 [38]. Coarse-grained and replica-exchange simulations yielded a structure for the TM domains with a stable interface. This model was strongly supported by its remarkable alignment with an independent AlphaFold2 prediction of Fzo1 in complex with its fusion partner Ugo1. Moreover, the pivotal role of Lys716 was validated, as its mutation disrupted mitochondrial fusion, and simulations demonstrated that Lys716 destabilized the membrane.

These works highlight the complexities involved in molecular modeling of mitofusins. As previously mentioned, predicting the three-dimensional structure of proteins has long been a fundamental and enduring challenge. Since proteins play a central role in biological processes, understanding their structure is essential to provide insights into their functions and mechanisms within cells, as well as into the designs of targeted therapies. Modeling proteins using bioinformatics tools is attractive as experimental methods can be time-consuming, expensive, and/or technically challenging. Prediction methods provide a faster and cost-effective means to predict 3D structures, which can be applied to all proteins, albeit with varying accuracy. In this context, AlphaFold 2 [39], AlphaFold 3 [40] and RoseTTaFold [41] are considered to provide significantly more reliable predictions than previous homology modeling approaches.

In this work, we explore the structure of mitofusins, particularly Fzo1 and Mfn1, using Alphafold. Full-length monomeric, dimeric, and tetrameric assemblies were generated, including complexes with proposed fusion partners such as Ugo1 and SLC25A46. Although AlphaFold yielded limited conformational variability for isolated monomers, greater structural diversity emerged upon modeling homo- and hetero-oligomeric states. Interactions between HR1 and HR2 domains of opposing mitofusins was predicted by AlphaFold, resulting in a dimer configuration in which the HR regions are not placed parallel to each other (as described in previous models [33, 34]), but instead interact in a crossed arrangement. This is described as a cross-type dimerization mode, a configuration not described in currently available experimental structures. The potential structural relevance of this interface is supported by recent experimental data. These full-length models enable us to formulate a hypothetical mechanism for outer mitochondrial membrane fusion. Together, these models offer a structural framework that may help refining current views of mitofusin-induced outer mitochondrial membrane fusion.

## 2 Materials and methods

### 2.1 Alphafold predictions

To predict the structures of mitofusins, we used Alphafold3 [40], and Colabfold 1.5.0 (AlphaFold2.3.1) [42] for custom sequence alignment predictions (see Supplementary Table **S2** and Figure **S7**). The various predictions are described in Supplementary Table **S1**.

#### Capri Evalutaion

Models with the best ipTM (or pTM for monomers) were subjected to a HAD-DOCK3 [43] energy minimization pipeline (*emref* module, default parameters, no restraints) and CAPRI evaluation (*caprieval* module, default paramters) to compare them to the corresponding experimental structures (Mfn1: 5YEW, 5GOF, 5GOM ; Mfn2: 6JFK ; Fzo1: 9KFE, 9KFF, 9KFD). The CAPRI evaluation relies on standard metrics such as interface and ligand RMSD (i-RMSD and l-RMSD), and the fraction of native contacts (FNAT), to quantify the accuracy of predicted protein–protein interfaces. To ensure a consistent comparison, predicted models were trimmed to match the sequence coverage of the corresponding experimental structures prior to evaluation. CAPRI-based evaluation was restricted to mitofusin components.

#### Custom Multiple Sequence Alignment

47 sequences of mitofusins were found using PSI-BLASTP version 2.13.0 [44, 45] and residues 702-760 of the Fzo1 sequence (Uniprot: P38297). The sequences obtained are listed in Supplementary Table **S2**, and the MSA of the TM domains is shown in Supplementary Figure **S7**. The full sequence of the mitofusins was then aligned using the T-coffee alignment procedure [46, 47]. This alignment was then provided to AlphaFold in A3M format for the prediction of the Fzo1 monomer.

#### Contact Analysis

Intermolecular contacts were analysed using the python library MDAnalysis [48, 49] with an inhouse script. The lowest distances between all residue pairs were measured using all the heavy atoms of the structures (e.g. except for hydrogen atoms, to stay coherent with the experimental structure).

### 2.2 Fzo1-Ugo1 oligomer simulations

A coarse-grained molecular dynamics simulation was performed, comprising the heterotetramer of 2 Fzo1 and 2 Ugo1 protein units built with Alphafold 3 [40].This setup was selected to investigate the stability and interaction patterns of a putative Fzo1–Ugo1 assembly.

#### System setup

For Fzo1, all produced AF models were refined using HADDOCK3 [43], after which the model with the best HADDOCK score was selected. This optimized model was then further processed using the CHARMM-GUI [50], adding a membrane consisting of 2000 lipids with a percentage composition of 45, 35, 14 and 6 for POPC, POPE, POPI and CD, respectively.

#### Simulation Parameters

The GROMACS 2022.5 program was used to perform the molecular dynamics (MD) simulation [51]. All molecular graphics were rendered using VMD [52]. The coarse-grained force-field MARTINI2 [53, 54] was used. For membrane-only and amphipathic helix systems, we followed the protocol of CHARMM-GUI which consisted in two energy minimizations of 5000 steps, followed by an equilibration of 5 simulations within the NPT ensemble. The velocity-rescaling thermostat [55] at 303.15 K, and the Berendsen barostat at 1 bar [56] were applied for a sequence of 5 simulations of 1 ns, 1 ns, 1 ns, 0.75 ns and 1 ns (with a timestep of 0.002 ps, 0.005 ps, 0.01 ps, 0.015 ps, 0.020 ps respectively). A production run of 10 *µ*s followed, at 310 K, using the velocity-rescaling thermostat [55] (lipid, water and proteins coupled separately) and 1 bar using the Parrinello-Rahman barostat [57] (compressibility of 3.0 *×* 10*^−^*^4^*bar^−^*^1^). Pressure coupling was applied semi-isotropically. A time step of 0.020 ps was used with the leapfrog integrator. Lennard-Jones interactions were cutoff at 1.1 nm. Bond lengths were constrained using the LINCS algorithm [58]. The reaction-field method [59] was used for evaluating electrostatic interactions, with a Coulomb distance cutoff of 1.1 nm, a relative dielectric constant of 15. The neighbor list was updated every 20 steps.

#### Simulation Analysis

Contacts between the two Fzo1 monomers were studied using the python library MDAnalysis [48, 49] with an in-house script.

## 3 Results and Discussion

### 3.1 AlphaFold exclusively produces monomeric mitofusin structures in an open conformation, incompatible with current hypotheses of membrane tethering mechanisms

The AF3 models of Mitofusin monomers using default settings are shown in Figure 2. First, all structures predicted in these conditions for the three molecules —Mfn1, Mfn2, and Fzo1— adopt an open conformation (Figures 2B,C,D and Supplementary Table **S1**). Figure 2 displays, on the left of each panel, the structures colored as a function of the pLDDT, and on the right, as a function of the domains (shown in Figure 2A). Each structure presents low pLDDT values in the TM domains (Figure 2B, C, D), and none show the expected two helix TM domain structure [38]. It is interesting to note that a kink is located at the conserved Arginine or Lysine residue (Figure 2B, C, and D). Furthermore, both Mfn2 and Fzo1 have low values in the N-ter section of the structure. In Fzo1, the N-ter section of the protein (before the HRN domain) is predicted as intrinsically disordered by AF.

**Figure 2:**
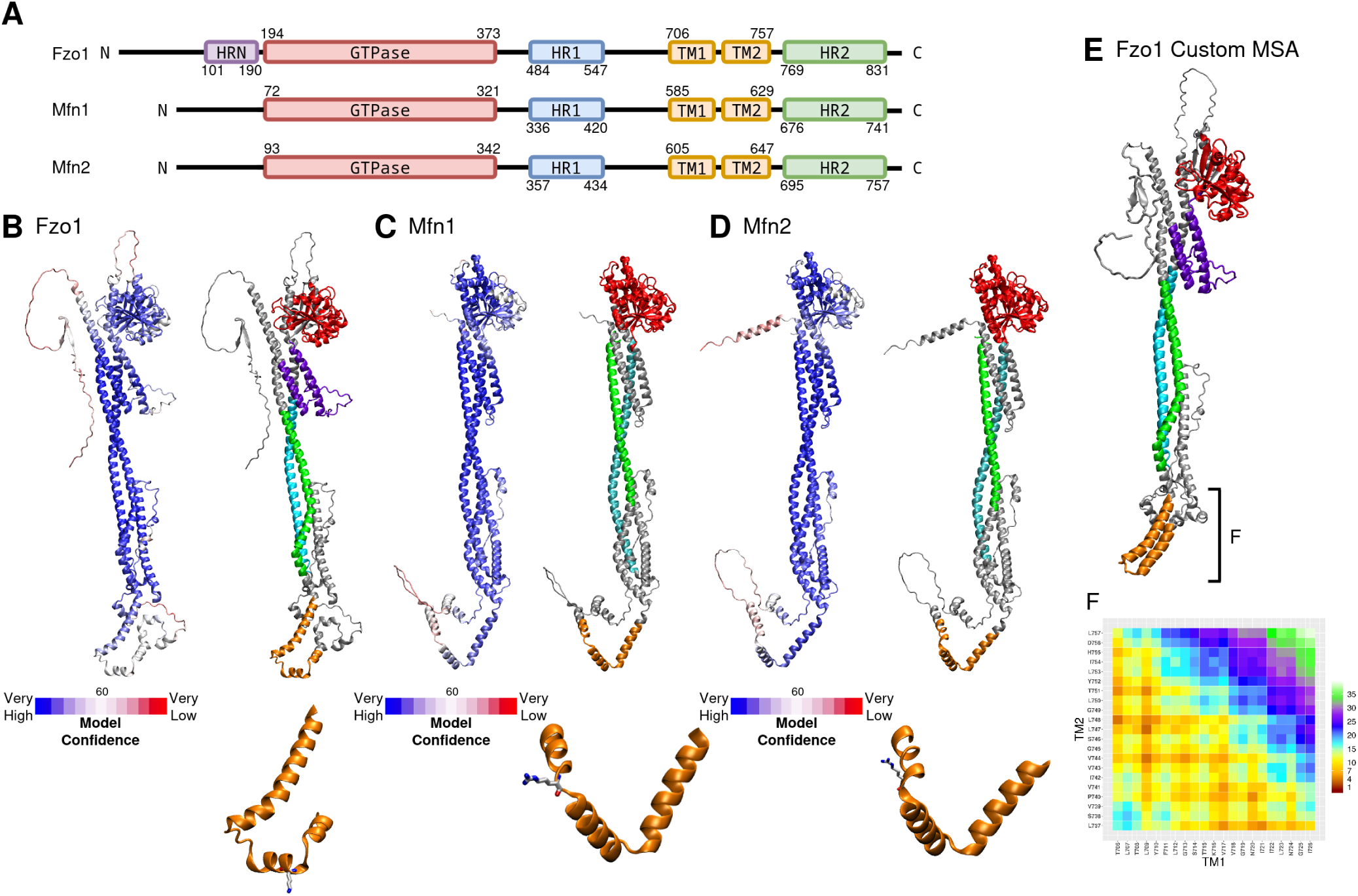
AlphaFold monomeric models of mitofusins. (A) Schematic representation showing the domain organization of mitofusins. GTPase domain in red; HR1/2, heptad repeat 1/2 in blue/green; HRN in purple; TM, transmembrane region in orange. Borders of each element are indicated by residue numbers. (B) Left, Fzo1 AlphaFold model colored as a function of pLDDT score. Right, Identical model colored as a function of the domains. The bottom is a zoom on the TM section of the protein, with the conserved Lys716 shown as licorice. (C) Left, Mfn1 AlphaFold model colored as a function of pLDDT score. Right, Identical model colored as a function of the domains. The bottom is a zoom on the TM section of the protein, with the conserved Arg594 shown as licorice. (D) Left, Mfn2 AlphaFold model colored as a function of pLDDT score. Right, Identical model colored as a function of the domains. The bottom is a zoom on the TM section of the protein, with the conserved Arg613 shown as licorice. (E) 2nd ranked model built using a custom MSA with AF2 and colored as a function of domains. The colors are described in (A). (F) Contact map between TM1 and TM2 of model (E). Each couple of residues are colored as a function of the distance (in Å).

The Fzo1 structure proposed by AF3 displays a very different structural organization compared to the models proposed in 2017 [34] and 2019 [35]. The HRN domain is folded beneath the GTPase domain. The placement is similar to the N-terminal helices observed in both Mfn1 and Mfn2 [6, 28]. Furthermore, the TM section exhibits significant differences, notably because AF3 predictions fail to generate the anticipated helical structures (Figure 2). The AF3 models are, however, not very consistent with the theoretical fusion processes previously proposed. The absence of a loop in the center of the mitofusins HR1/2 coiled-coil implies strong conformational rearrangement to fit the hypothesized mechanisms described in 2017 [34]. To be in accordance with previously proposed mechanisms (specifically presented in 2018 [60] and 2017 and 2019 [34, 61]), the helix bundles of the mitofusin models would have to locally break and unfold into a loop. However, none of these fusion models have been validated by experimental data. Therefore, new mechanisms can be proposed from the analysis of the models produced by AF, and more specifically from the analysis of multimers of mitofusins.

#### 3.1.1 Custom MSA provides additional insights into TM domain interactions and structure prediction

The main limitation of the Fzo1 AF3 model is its transmembrane (TM) section, where the low pLDDT values and the consistent presence of a kink in the TM1 helix indicate lower confidence in this region. We therefore tested the use of a custom MSA with AF2, following recent work showing that tailored MSAs can influence AlphaFold predictions by biasing conformational sampling toward specific structural features [62, 63]. Figure 2E shows the model produced using an MSA composed of 48 sequences of fungal species (see Supplementary Table **S2**). This MSA is particularly well aligned on the TM domain of Fzo1 (Supplementary Figure **S7**). The TM domains (TM1 and TM2) were predicted to interact with each other. Although the HR and GTPase domains show good alignment, the positions of the two transmembrane helices differ compared to those in the previously presented models (Figure 2B–D), as shown in Figure 2F. The residues involved in interactions are similar to the ones described previously in 2024 [38]. These interactions are shown for the best ranked model according to ipTM (0.54) in Figure 2E. All conformations show the same residues of the helices involved in the interface.

From these observations we can conclude that, when AF is given a more specific alignment, it converges to a two helices type of TM domain (with interactions between the two helices), consistent with our proposed TM domain model [38]. These observations are consistent with recent work showing that AlphaFold predictions are highly sensitive to MSA composition [62, 63], and that tailoring evolutionary input can bias models toward specific structural solutions. However, as in our case, such approaches primarily improve local structural features and convergence, while large-scale conformational transitions remain difficult to capture.

### 3.2 Oligomers involving solute carriers shed new light on cross-type cis-dimers

While monomeric models provide useful insights into the overall architecture of mitofusins, mitochondrial fusion is widely thought to rely on the formation of dimers and oligomers of mitofusins [64, 60, 35, 65, 29]. In this context, we extended our analysis to oligomeric assemblies of mitofusins, including dimers and tetramers, both in isolation and in the presence of proposed interaction partners, the solute carriers Ugo1 or SLC25A46. This approach allows us to explore how intermolecular interactions may influence mitofusin conformation and to identify recurring structural features that could help us propose a new fusion mechanism.

#### 3.2.1 Mitofusin dimer predictions are influenced by the presence of solute carriers

Tetramers involving two mitofusins and two solute carriers (Ugo1 for *S. Cerevisae* and SLC25A46 for humans) yielded the most interesting results. Their models are shown in Figure 3 (mitofusins are shown alone) and Supplementary Figure **S8** (complete models). The various oligomers predicted for this study have been listed in Supplementary Table **S1**. Fatty acids were used in some of our predictions, as it was previously shown to improve predictions [66]. First and foremost, a novel dimerization pattern for mitofusins emerges: cross-interactions involving both the GTPase domain and HR coiled-coil are prevalent in most models containing two mitofusins. Closest contacts between mitofusins are shown in Supplementary Table **S3**. It can be observed that the main contacts observed are located outside of the TM domains. Furthermore, the dimeric structure seems to be maintained through salt bridges, observed in the HRN (for Fzo1 only), GTPase, and HR2 domains. Indeed, polar and charged residues are able to form salt bridges, and hydrogen bonds are observed in close proximity for both Mfn1 and Fzo1.

**Figure 3:**
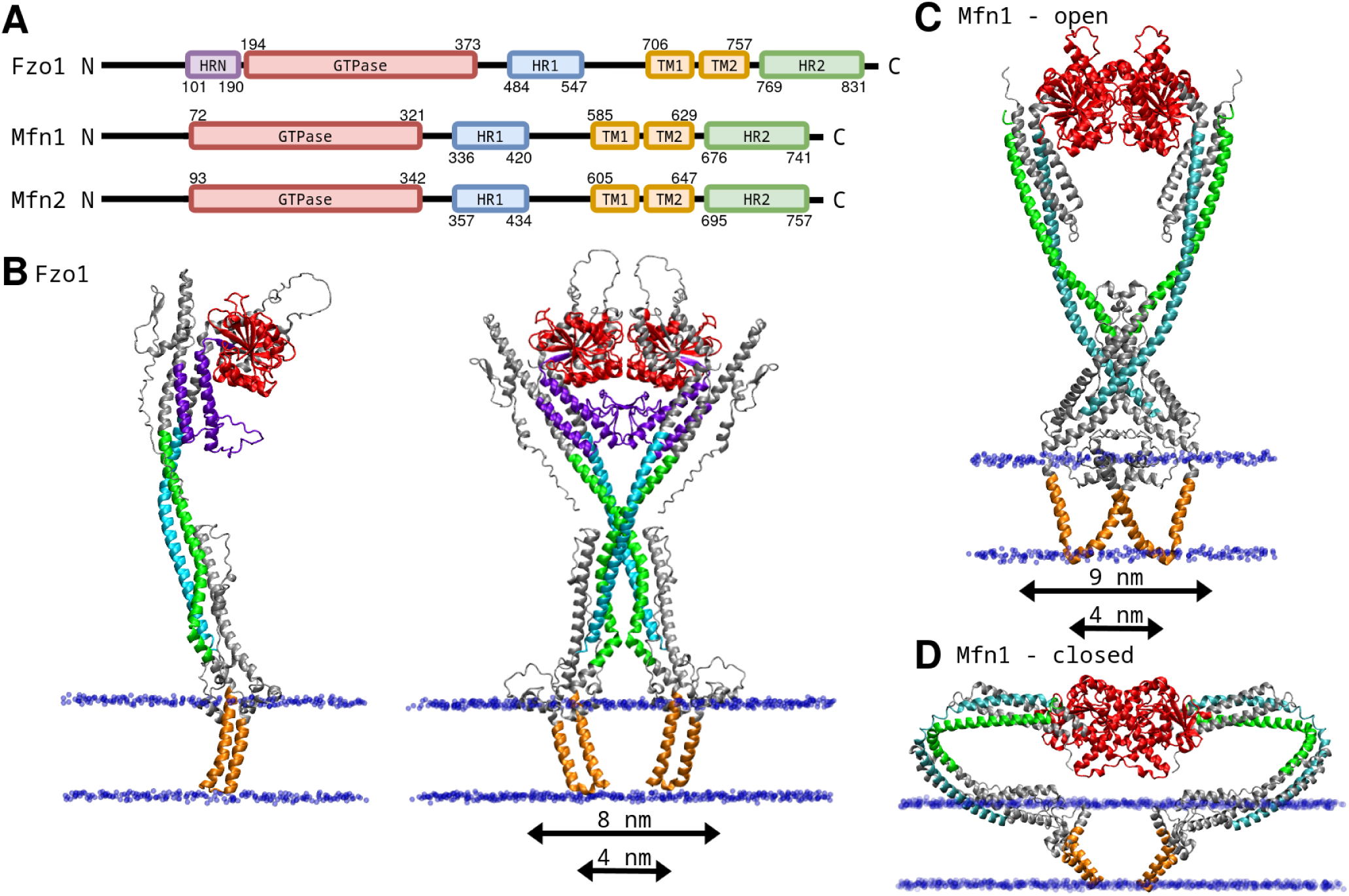
Oligomeric models of mitofusins. (A) Schematic representation showing the domain organization of the mitofusin Fzo1. GTPase domain in red; HR1/2, heptad repeat 1/2 in blue/green; HRN in purple; TM, transmembrane region (including the loop in between TM1 and TM2) in orange. Boundaries of each element are indicated by residue numbers. (B) Monomer and dimer of Fzo1 produced by AF with two Fzo1 and two Ugo1 originally in the system (see Supplementary Figure **S8**A). The domains are colored as a function of the domains defined in (A). The unassigned sections of the protein are colored in grey. The phosphate atoms of the membrane are represented in blue van der Waals representation. The dimer spans approximately 8 nm at the level of the GTPase domains (10 nm including the helices) and narrows to about 4 nm at the HR domains. (C,D) Dimers of Mfn1 produced by AF with two Mfn1 and two SLC25A46 (see Supplementary Figure **S8**B). Open cis-dimer on top (C), closed cis-dimer on the bottom (D). The domains are colored as a function of the domains defined in (A). The unassigned sections of the protein are colored in grey. The phosphate atoms of the membrane are represented in blue van der Waals representation. The HR domains maintain their helical structure throughout the sequence. The open dimer (C) spans approximately 9 nm at the level of the GTPase domains (helices included) and narrows to about 4 nm at the HR domains.

The dimerization mode is very different from the BDLP homolog [32] as well as the Fzo1 models proposed in 2019 [35]. Therefore, a molecular dynamics simulation of the system was performed (Supplementary Figure **S8**A) to assess the stability of the protein–protein interactions outlined above. Fzo1 monomers maintained contacts and close interactions throughout the production run (Supplementary Table **S3**), and a number of the residues forming salt bridges identified between the Fzo1 monomers were found in close proximity throughout the simulation.

When GTP was added to the AF predictions, it was found as expected in the GTPase domain (Supplementary Figure **S8**E). However the absence or presence of GTP/GDP was not found to impact significantly the models. In fact, this observation is coherent with the work of Škrinjar et al. [67] who showed that nucleotide binding alone does not necessarily impose a large-scale conformational rearrangement in protein structures. In light of these findings, and given that our predictions did not reveal nucleotide-dependent differences at the global level, we did not further investigate the impact of GTP/GDP binding in this study. Instead, we focused on oligomerization modes and membrane-related configurations, which appear to play a more decisive role in shaping mitofusin conformations.

Most models involving two mitofusins and two solute carriers present open conformations of mitofusins. Interestingly, models involving two human mitofusins Mfn1 and two SLC25A46 show a diversity of conformations for the mitofusins involved. Figure 3D shows a closed conformation of Mfn1. Other conformations obtained for this system are shown in Supplementary Figure **S10**A-E. Some of the models presented in the latter figure are however quite unrealistic, as they show the coiled-coil globular portion of the protein spanning through the membrane plane (Supplementary Figure **S10**A, E). The HR domains are shown bent, and the secondary structures appear mainly conserved, which differs from the BDLP homolog [30, 32], in which an unstructured loop section was observed. AF seems to show here potential flexibility in the coiled-coil section of mitofusins. Kinks can be observed in the bent coiled-coil of the model presented in Figure 3D and Supplementary Figure **S8**B. These cross-interactions involving the coiled-coil portions of the proteins contrast with previsouly predicted dimers of Fzo1 [35], all based on the homologue BDLP [30, 32]. Brandner and collaborators produced in 2019 [35, 31] a dimerization only involving the GTPase domain. At the time, this was consistent with the BDLP structure, in which the interaction interface also primarily involves the GTPase domain [32].These interactions can be observed when no lipids or solute carriers are placed in the system, as seen for Mfn2 in Supplementary Figure **S8**D.

Furthermore, Ugo1 and SLC25A46 (both tranmembrane proteins) are localized on the membrane plane, i.e. the same level as the TM domain of the mitofusins involved (shown in Supplementary Figure **S8**A and B). Consequently, the structure of the TM domain of mitofusins is impacted. In the absence of solute carriers, the transmembrane helices (TM1 and TM2) of mitofusins appear separated from each other and show kinked helices (Figure 2 and Supplementary Figure **S8**C, D, E). With the presence of Ugo1/SLC25A46, a more compact conformation of the domain is evident (Figure 3B, C, D). These models were built in the presence of fatty acids, which in the predicted models localize where the membrane would be (Supplementary Table **S1** and Supplementary Figure **S8**E). In Fzo1, the structure of the TM domain and the interactions between the helices previously described in Figure 2 and in 2024 [38] is observed in the presence of Ugo1. However, none of the human mitofusin models presented here show close interactions between the helices TM1 and TM2 (Figure 3C and Supplementary Figure **S4**).

Finally, it can be noticed that Ugo1 N-termini are located in the intermembrane space, which is unexpected, as the opposite was described by Sesaki et al. in 2001 [21]. However, the C-term is on the IMS side which is consistent with experimental data [21]. The second extended intrinsically disordered domain of the protein is located on the cytosol side of the membrane, and interacts with the HR domains. Similarly, the N-termini disordered sections of SLC25A46 are interacting with the human mitofusins’ coiled sections. In fact, some of the closest contacts between the mitofusins and their solute carriers involve the intrinsically disordered sections of those solute carriers (described in Supplementary Table **S4**).

#### 3.2.2 The AlphaFold oligomers are in agreement with the available experimental data

Interactions between the mitofusin monomers were analysed and are shown in Supplementary Table **S3**. As shown by the grey rows of the tables, the closest inter-chain contacts correspond to contacts observed in the experimental structures [6, 7, 28]. Numerous contacts at a distance lower than 3Å are found between monomers of the Mfn1 open conformation model (Figure 3C). The interactions identified within the region of the model corresponding to the experimentally solved structure 5YEW [7] overlap with those reported in the crystal structure (see Supplementary Table **S3**B). For the closed conformation of Mfn1, the partial structure 5GOM [6] is used for comparison purposes. Here, the closest contacts are found within the GTPase domain, many of which are also observed in the experimental structure (Supplementary Table **S3**C). The alignment of these AF models with experimental structures are shown in Supplementary Figure **S9**B. The RMSD between AF and the experimental structures is low (Supplementary Figure **S9**). The ‘caprieval‘ module of HADDOCK3 [43] evaluates open and closed conformation structures as medium quality models (Supplementary Figure **S9**B). This result is expected for Mfn1, as these structures (PDBs: 5GOM[6], 5YEW[7], 5GOF[6], 6JFK[28]) were released before 30th of September 2021 and are part of AlphaFold 3 training data.

On the other hand, Fzo1’s structure was not available to Alphafold at the time of making the models presented here. First, in a previously proposed Fzo1 model [34, 35] various salt-bridges and hydrophobic interactions were found, and were experimentally validated using swap mutations[34]. These salt bridges are described in Supplementary Figure **S9**A. A first salt-bridge was found involving the Lys464 residue, a target for post-translational modification by ubiquitin in Fzo1 [33], and Asp335. Single point mutations D335K and K464D induced a total inhibition of respiratory growth, while the swap mutation D335K-K464D partially but significantly corrected the defect. Similarly, the K200-D313 salt-bridge and the D523-H780 hydrogen bond were found, although the H780D mutant did not show respiratory inhibition [34]. Supplementary Figure **S9**A underlines that salt bridges D335K-K464D and K200-D313 predicted in 2017 were found in our new AF3 Fzo1 dimer models (and validated in the experimental structure, result not shown), while D523-H780 residues are found to be quite far apart in our new model. However, the D523-H780 salt-bridge could be reasonably questioned, as one of the two single point mutants does not show respiratory inhibition. Hydrophobic contacts were hypothesized as well in the 2017 study [34]. L501-L504-L802 and T490-L819 were observed to be in close contact, yet they are farther apart in our new AF3 model.

Recently, Yan et al. reported a new structure [29], providing additional insights that were not accessible from earlier experimental data and that allow discrimination between our current model and the 2017 model [34]. While both AlphaFold2 [39] and AlphaFold3 [40] were not trained on this structure, both tools were able to produce a similar dimerization mode, as observed by Versini et al. in 2023 [68]. In this work, we show the two aligned structures in Supplementary Figure **S9** as well as the common contacts found in the experimental structure, and the AF3 model in Supplementary Table **S4**. Most of the closest contacts found in the AF3 model are found in the resolved part of the experimental structure. The AF3 model reproduces more than 75% of the correct contacts between the monomers. Alltogether, these results strengthen the validity of the AF3-derived models and support the relevance of the proposed cross-type dimerization mode.

#### 3.2.3 Structural hypotheses on mitofusin tetramers

When looking at mitofusins and solute carrier models (Figures 3), it appears that all models can be categorized as cis-dimer, i.e. both monomers would be located on the same membrane. These models could represent either the initial or final conformations of mitofusins in the membrane fusion process. However, the limited experimental evidence available makes it difficult to draw definitive conclusions. On the other hand, homotetramers of Mfn1 show high conformational diversity, as seen in Figure 4. Notably, the models consistently display interactions between GTPase domains, and proximity between the HR domains is often observed as well. The mode of tetramerization varies between the different predictions, with the interfaces forming at distinct regions of the HR domains. Furthermore, the models in Figure 4A, B and C are most likely trans-tetramer models of Mfn1, while D and E show cis-tetramer models of respectively Mfn1 and Fzo1. In the case of models B and C, the parallel membranes could be positioned in two alternative ways, described in the figure. Furthermore, the cyan membranes in Figure 4B, are at a distance of about 9nm,

**Figure 4:**
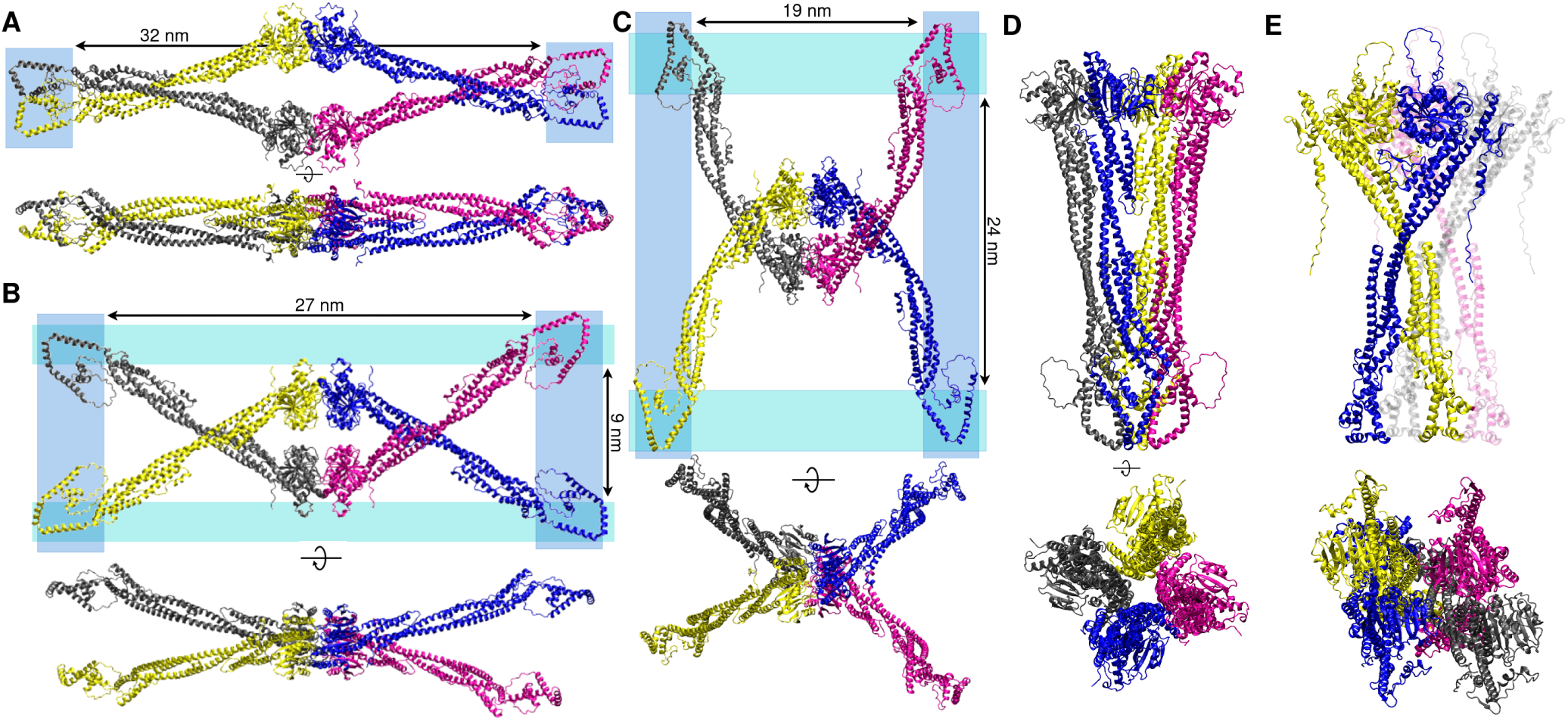
Tetramer models of mitofusins. (A),(B),(C),(D) Cartoon representation of Mfn1 tetramers. Monomers are represented in a different color for readability. Blue or cyan rectangles are positioned at potential membrane positions. (E) Cartoon representation of Fzo1 tetramers. (A),(B),(C) are trans tetramers. (D),(E) are cis-tetramers.

Tetramer models involving fatty acids, as presented in Supplementary Figure **S10**F, G show only open cis tetramers. AF3 failed to produce Fzo1 tetramer, as shown in Supplementary Figure **S10**G in which two dimers are superimposed. The predicted TM domains were partially disordered and kinked, seemingly interacting with the surface of the membrane environment built by the fatty acids. This is in direct opposition to previous studies outlining the position of TM domains [69, 64, 70]. Hence, these models are currently disregarded.

As we are unable to discriminate between the different models in Figure 4 without additional experimental data, no definitive conclusion can yet be drawn regarding the precise oligomerization mode of mitofusins. Furthermore, experimental analyses would be essential to validate or refute the alternative assemblies proposed here. However, each of these models could be considered as potential alternative conformations that may coexist under different conditions or at different point of the fusion process. Individual models presented in this work could be compatible with a membrane tethering model. However, at this stage, the information provided in this work is hard to integrate into a single, unified mechanism of fusion. Nonetheless, these trans-tetramers may inspire a novel structural basis for bringing opposing membranes into close apposition.

### 3.3 What about the mitofusin-mediated mitochondrial fusion mechanism ?

The overall fusion mechanism of mitofusins remains elusive. Brandt et al. 2016 [61] have detailed various conformations of the outer mitochondrial membrane when fusion is proceeding, showing a tethering and docking ring stage before fusion. The OMs’ tethered intermediates were observed to be at a distance of about 6 to 8 nm. Macromolecular assembly was observed within the ring-shaped mitochondrial docking complex, which was hypothesized to be mitofusins. Based on these observations and the structural analysis of mitofusins, many hypotheses on the mitochondrial fusion process were formed. As mentionned before, the GTPase domain is known to drive dimerization [32], while the heptad repeat domains were shown to participate in membrane tethering as well as membrane destabilization [64, 60, 65]. The fusion process is often thought to be involving mitofusins acting in trans configuration to tether together adjacent mitochondrial target membranes. Daste et al. 2018 [60] and Cohen et al. 2018 [65], proposed a mechanism involving the human mitofusin Mfn1, based on the structures solved in 2017 and 2018 [6, 7]. A conformational rearrangement would occur subsequent to trans-dimerization through the GTPase domain, which would position HR1 in a way that can destabilize the membrane. In 2019 [35] membrane tethering models based on cis- and trans-oligomers were investigated, taking advantage of the Fzo1 model produced in 2017 [34], in its closed conformation. In this proposed model, two Fzo1 molecules would form oligomers in cis configuration *via* their GTPase domain, and hydrophobic interactions would be formed between Fzo1 in trans. Consequently, the spines of HRs would align in an antiparallel manner, consistent with the crystal structure of Mfn1 [64].

Recently, a new working model for yeast OMM fusion was proposed based on the latest experimental partial structures of Fzo1 [29]. In this study, the authors integrate structural and biochemical data to describe a fusion mechanism in which Fzo1 molecules initially form trans dimers upon GTP loading, tethering opposing mitochondrial membranes. Subsequent GTP hydrolysis drives a conformational transition that brings the membranes into close proximity and promotes fusion.

In this work, we described both cis and trans oligomers. These oligomers are however quite different from the models previously established. Based on our models of mitofusin dimers, it is tempting to propose a mechanism (Figure 5). As a matter of fact, the cross-type cis-dimers could be the result of mitofusin induced membrane fusion. A first interaction between the GTPase domains of trans mitofusins would occur (as seen in Mfn1 dimer of Supplementary Figure **S8**F and Mfn1 trans-tetramers of Figure 4). The HR interactions would then be induced, leading to the cross-type interactions observed in Figure 3B-C. These interactions lead to dimers spanning between 4 nm to 8 nm, which would be in accordance with Brandt et al., who reported that the membranes where at 6-9 nm appart at the edges of the contact sites [61]. In this final step completing membrane tethering and leading to fusion, the GTP has been hydrolysed into GDP, as it was revealed to be necessary for membrane docking and fusion by Brandt et al. [61]. Post-fusion cis-dimers are the result of membrane fusion, observed in Figures 3B-C. The hypothetical fusion mechanism is shown in Figure 5. Unlike the recently proposed Fzo1-mediated fusion model [29], our mechanism explicitly incorporates cross-type interactions between mitofusins, which emerge strongly from our structural models.

**Figure 5:**
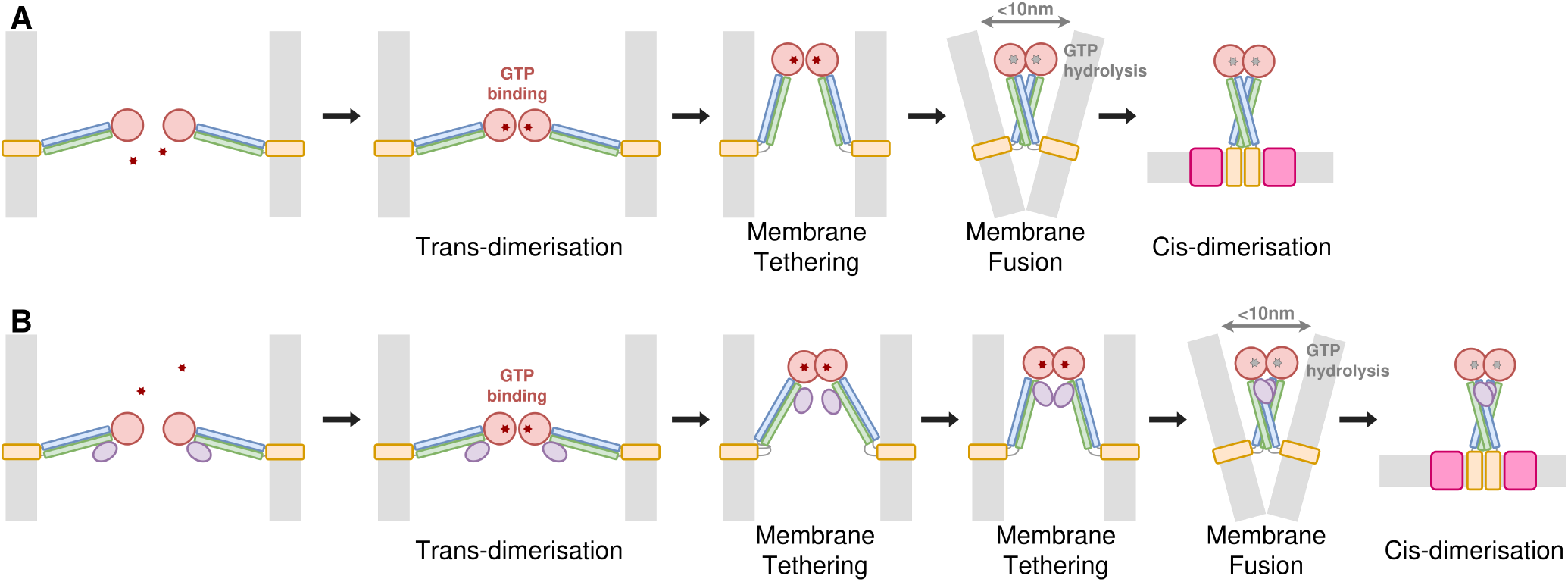
Proposition of a Mitofusin induced membrane fusion mechanism based on AF models and available experimental data. Mfn1 (A) and Fzo1 (B) are schematized, with the main domains represented as rectangle or circles: the orange boxes represent the TM domain, the blue and green long rectangle represent respectively HR1 and HR2 domains, the red circle is the GTPase domain and the purple circle is the HRN domain. The solute carrier partner (either Ugo1 or SLC25A46) is represented as a pink square. In light grey stripes are represented the membranes. The red stars are GTP molecules, and the grey stars are GDP. The first step is the trans-dimerization of mitofusins through the GTPase domain upon GTP binding. Then is followed conformational changes leading to membrane tethering. Other conformational changes lead then to cross-dimers, and membrane fusion. In the final step proposed, the initial trans-dimer became a post-fusion cis-dimer, stabilized by the solute carrier, either SLC25A46 (A) or Ugo1 (B).

Other important conformational changes could be required to facilitate the outer mitochondrial membrane fusion, as expected from members of the dynamin superfamily [32, 71, 72, 73]. Our current proposed fusion mechanism does not involve the bending capabilities of the HR domains observed in the closed conformation of Mfn1 (Figure 3D). While the rupture of coiled-coil structures might be conceivable and consistent with previously proposed mechanisms of the yeast mitofusin [34, 35] and BDLP [30, 32], these structures could also undergo bending without necessitating helical breakage, bringing the two opposing membranes closer together and possibly interacting with the membrane, upon reorganization of the HR domain hydrophobic spines.

We were not able to include the tetrameric conformations in the fusion mechanism, as their structural diversity and the lack of experimental validation currently prevent a clear interpretation of their functional relevance. Our model however includes this new cross-type dimerization, not reported before in previous work. Furthermore, by reconciling structural predictions with available experimental data, our work proposes a hypothesis for the fusion mechanism, that could guide future biochemical and structural experimental investigations.

## 4 Conclusion

In this work, the modeling of mitofusins has been undertaken using AlphaFold complemented with HADDOCK and molecular dynamics. Monomers, dimers and tetramers involving fusion partners Ugo1/SLC25A46 were produced and analysed. While AlphaFold generally provides limited conformational diversity for an isolated mitofusin monomer, structural variability emerged when the molecular environment was modified through oligomerization or inclusion of partner proteins. The presence or absence of GTP did not produce a detectable global structural impact on our predictions. In contrast, clear structural differences were observed upon dimerization of mitofusins, as well as upon association with solute carriers and membrane-binding partners—both individually and in combination. In this context, AlphaFold predicted a previously undescribed cross-type dimerization mode involving interactions between the HR domains, a configuration not reported in the current literature.

Recent AlphaFold Database–derived models of Fzo1 have proposed dimeric assemblies of the protein [74]. However, the prediction was based on a truncated constructs lacking the N-terminal region, including the HRN domain. This domain was shown to be involved in the interactions in the recent experimental structure of Fzo1 [29]. Hence, their partial nature limits their applicability to full-length mitofusin organization. In this context, our full-length modeling approach enables the exploration of oligomerization modes that involve the complete protein architecture.

It is worth noting, however, that most of the AlphaFold models presented in this work show low pTM/iPTM scores (see Supplementary Table **S1** and Figure **S6**). Many observations are however supported by experimental data. Indeed, previous work indicated that the mitofusins dimerize [6, 7, 29], as well as interact with their associated solute carriers [21, 27, 75, 76, 77, 25, 26]. These experimental data give some validity to the AF models despite their low confidence scores.

Most models showed mitofusins in an open extended conformation, and an absence of hinges in coiled-coil areas, in opposition to the bacterial homologue BDLP [30, 32]. Moreover, the cross-type cis-dimerization observed in dimers and tetramers of mitofusins contrasts with the bacterial homologues, as this type of dimerization was not observed when BDLP dimer form was solved [32]. This might be attributed to the relatively low similarity (about 43% [34]) between BDLP and mitofusins, leading to structural differences. Interestingly, the structures involving the mitochondrial solute carriers SLC25A36 (humans) showed that these inter-protein HR interactions are possible between any combination of mitofusins. Furthermore, AlphaFold was able to place both solute carriers, Ugo1 and SLC25A46, and fatty acids at the same location as the transmembrane helices of mitofusins, improving the helical structures of this region and finding interactions between helices of the same monomer previously reported [38].

A limitation of the recent experimental studies is the lack of structural information for the helix bundle and transmembrane domains. These regions are likely to contribute to mitofusin oligomerization and OMM fusion. By explicitly modeling TM-containing oligomeric assemblies and identifying cross-type cis and trans interactions, our study provides structural hypotheses for how these poorly characterized domains may participate in the fusion process. In this context, our models complement existing experimentally derived mechanisms by proposing additional modes of mitofusin assembly that could operate during or after membrane merging.

We concluded our work by proposing an OMM fusion mechanism for both mammals (Mfn1) and yeast (Fzo1). This fusion mechanism requires conformational changes in order to drive the membrane fusion, transitioning from trans-dimers on opposite membranes to cross-type cis-dimers. Future works could explore the mitofusin conformational space further, with tools such as AlphaFlow [78] or BioEmu [79], which are designed to explore a broader range of protein conformations. Given the computational nature of this study and the limited experimental validation available for full-length mitofusins, the proposed structural arrangements and fusion mechanism should be considered as hypotheses that require further experimental investigation.

## Appendix

**Supplementary Table S1:**
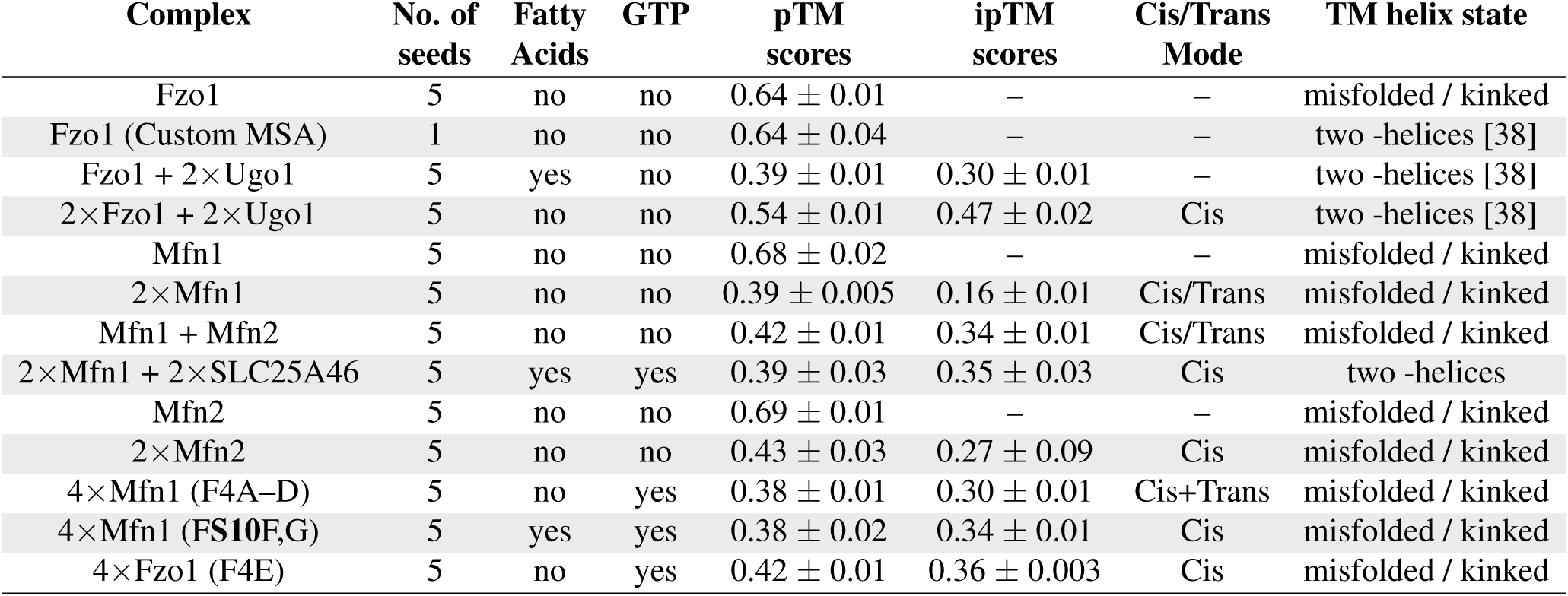
Summary table of the models produced with AlphaFold. The provided sequences were taken from UniProt [80] for Fzo1 (P38297), Mfn1 (Q8IWA4), Mfn2 (O95140), Ugo1 (Q03327), and SLC25A46 (Q96AG3). Fatty acids correspond to a mixture of oleic and palmitic acids. The “TM helix state” column provides a standardized qualitative assessment of helix folding and organization. Each seed generates five models. pTM and ipTM correspond to the AlphaFold predicted TM-score and interface TM-score, respectively. One seed produces to 5 models.

| Complex | No. of seeds | Fatty Acids | GTP | pTM scores | ipTM scores | Cis/Trans Mode | TM helix state |
| --- | --- | --- | --- | --- | --- | --- | --- |
| Fzo1 | 5 | no | no | $0.64 \pm 0.01$ | – | – | misfolded / kinked |
| Fzo1 (Custom MSA) | 1 | no | no | $0.64 \pm 0.04$ | – | – | two -helices [38] |
| Fzo1 + 2×Ugo1 | 5 | yes | no | $0.39 \pm 0.01$ | $0.30 \pm 0.01$ | – | two -helices [38] |
| 2×Fzo1 + 2×Ugo1 | 5 | no | no | $0.54 \pm 0.01$ | $0.47 \pm 0.02$ | Cis | two -helices [38] |
| Mfn1 | 5 | no | no | $0.68 \pm 0.02$ | – | – | misfolded / kinked |
| 2×Mfn1 | 5 | no | no | $0.39 \pm 0.005$ | $0.16 \pm 0.01$ | Cis/Trans | misfolded / kinked |
| Mfn1 + Mfn2 | 5 | no | no | $0.42 \pm 0.01$ | $0.34 \pm 0.01$ | Cis/Trans | misfolded / kinked |
| 2×Mfn1 + 2×SLC25A46 | 5 | yes | yes | $0.39 \pm 0.03$ | $0.35 \pm 0.03$ | Cis | two -helices |
| Mfn2 | 5 | no | no | $0.69 \pm 0.01$ | – | – | misfolded / kinked |
| 2×Mfn2 | 5 | no | no | $0.43 \pm 0.03$ | $0.27 \pm 0.09$ | Cis | misfolded / kinked |
| 4×Mfn1 (F4A–D) | 5 | no | yes | $0.38 \pm 0.01$ | $0.30 \pm 0.01$ | Cis+Trans | misfolded / kinked |
| 4×Mfn1 (FS10F,G) | 5 | yes | yes | $0.38 \pm 0.02$ | $0.34 \pm 0.01$ | Cis | misfolded / kinked |
| 4×Fzo1 (F4E) | 5 | no | yes | $0.42 \pm 0.01$ | $0.36 \pm 0.003$ | Cis | misfolded / kinked |

**Supplementary Figure S6:**
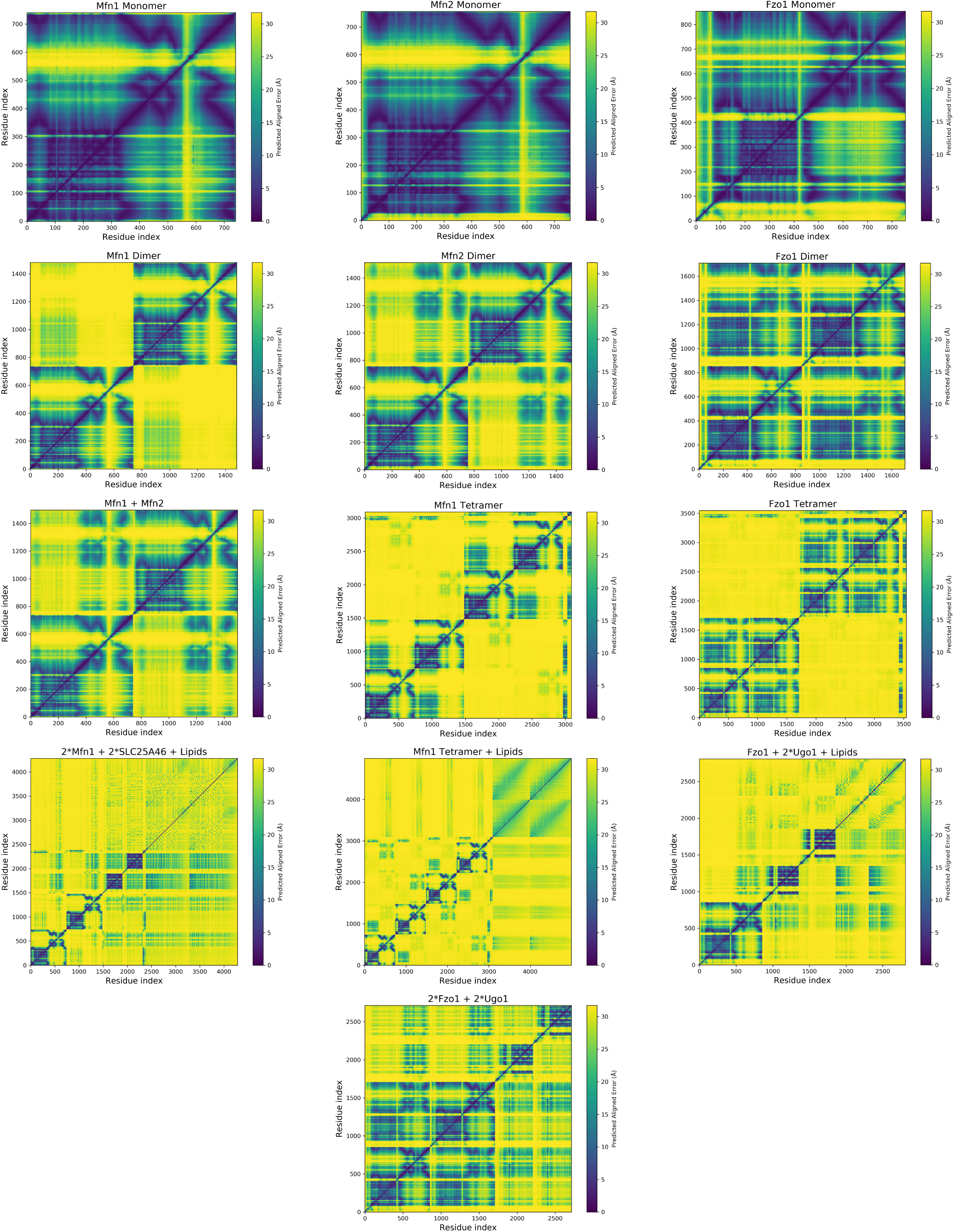
Predicted Aligned Error (PAE) matrices for the main models considered in this study. The selected PAE matrices were extracted from the top-ranked models based on the highest ipTM scores (or pTM scores in the case of monomers).

**Supplementary Table S2:**
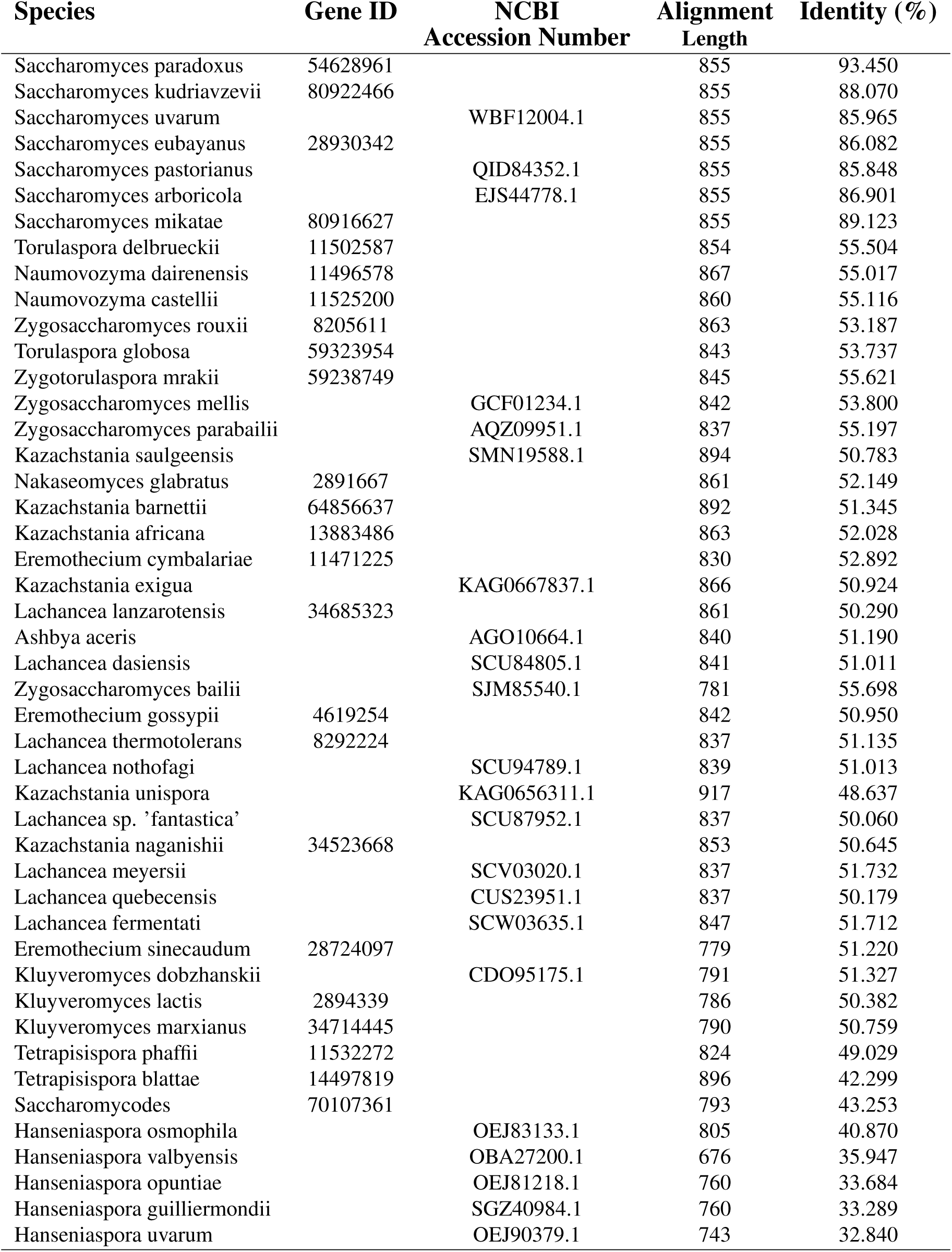
Summary table of the sequences selected for a custom alignment. When Gene ID could not be retrieved, the NCBI Accession Number was writen instead. Identity stands for amino acid identity. The percentage of identity with the mitofosin Fzo1 of Saccharomyces Cerevisae is written in the last column.

**Supplementary Figure S7:**
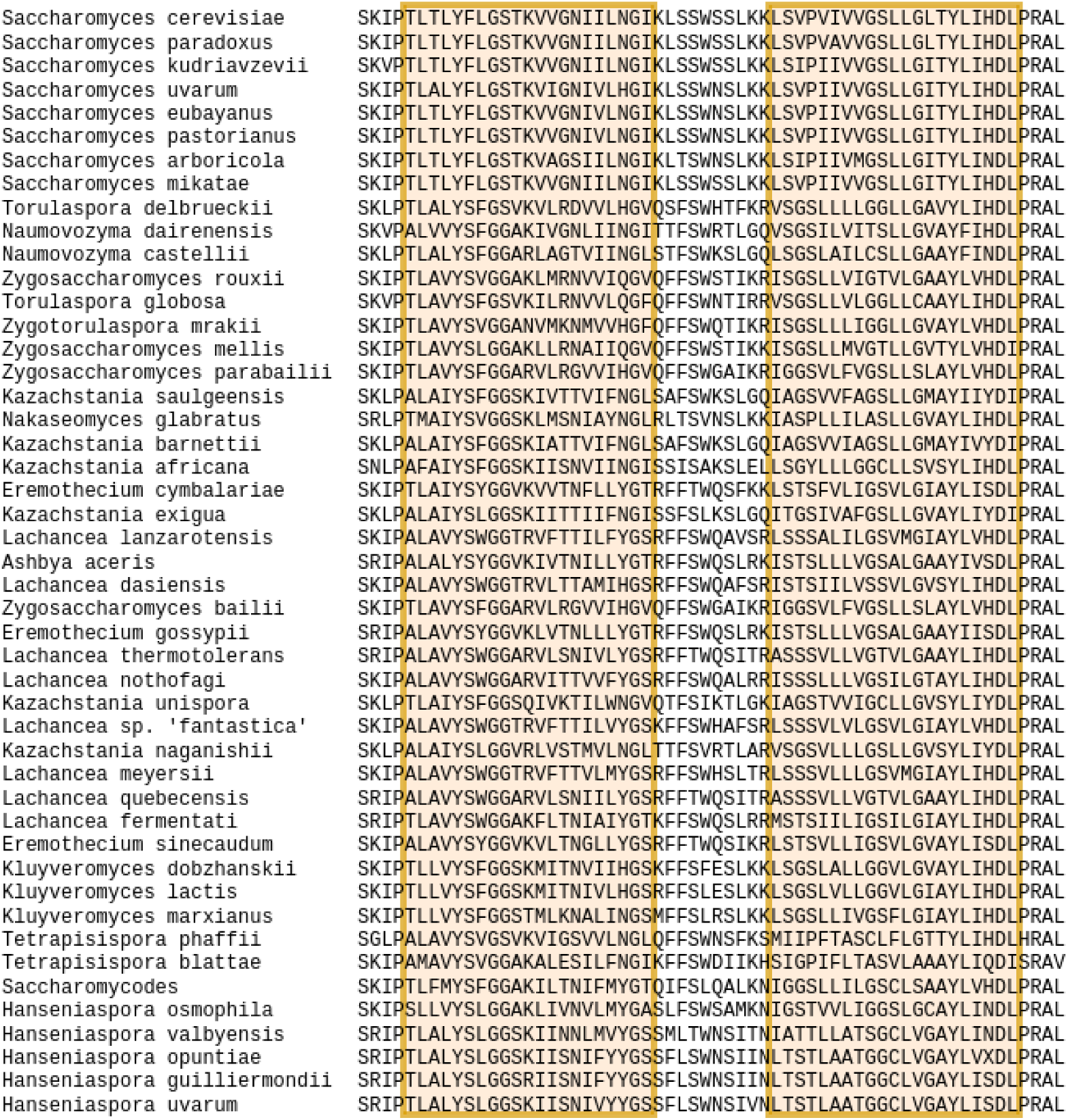
TM domain alignment of the mitofusins used for the custom MSA. Here is the TM domain alignment of the custom MSA. Each predicted TM helix is represented as a orange rectangle.

**Supplementary Table S3:** Distances in Å between the closest residues of mitofusin monomers in mitofusins dimers. (A) Residues at a distance lower than 3Å of the Fzo1 homodimer AF model. The model used for this analysis is shown in Figure 3B and Supplementary Figure **S8**A. The prevalence observed in the coarse-grained (CG) simulations is reported in the last column. This value was calculated as the fraction of simulation time during which the residues were found to be in contact. In the simulation, residues separated by a distance of less than 7Å were considered in contact, corresponding to the cutoff strictly used in the coarse-grained representation of the system. (B) Residues at a distance lower than 3Å of the Mfn1 open cis-homodimer AF model. (C) Residues at a distance lower than 3Å of the Mfn1 closed cis-homodimer AF model. For all (A), (B) and (C), bolded residues represent salt bridges. The models are compared to the experimental structure 9KFD [29] for (A) (note: LYS149 was not solved for chain B, i.e. not in the experimental stucture), 5YEW [7] for (B) and 5GOM [6] for (C).

| A | B | Distance<br>model (Å) | Distance<br>exp (Å) |
| --- | --- | --- | --- |
| THR84 | SER210 | 2.83 | 2.72 |
| ILE105 | GLU245 | 2.71 | 3.06 |
| GLY106 | GLU209 | 2.74 | 2.95 |
| HIS107 | GLN255 | 2.94 | 2.65 |
| ASN140 | GLU247 | 2.99 | NA |
| HIS147 | ASP251 | 2.97 | 6.46 |
| ASP183 | ASN214 | 2.79 | 2.91 |
| GLU209 | GLY106 | 2.8 | 2.91 |
| **GLU209** | **ARG238** | 2.61 | 3.45 |
| SER210 | THR84 | 2.81 | 2.79 |
| ASN214 | ASP183 | 2.86 | 3.0 |
| **ARG238** | **GLU209** | 2.64 | 3.43 |
| **GLU245** | **LYS99** | 2.49 | 5.36 |
| TYR248 | HIS144 | 2.64 | 3.92 |
| ASP251 | HIS147 | 2.85 | 6.4 |
| GLN255 | HIS107 | 2.96 | 2.62 |
| **GLU403** | **LYS511** | 2.65 | NA |
| ASN506 | HIS667 | 2.71 | NA |
| **LYS511** | **GLU403** | 2.55 | NA |
| THR562 | GLU652 | 2.74 | NA |
| PRO563 | ARG655 | 2.78 | NA |
| THR564 | ARG655 | 2.75 | NA |
| GLU652 | THR562 | 2.7 | NA |
| ARG655 | PRO563 | 2.88 | NA |
| ARG655 | THR564 | 2.71 | NA |
| HIS667 | ASN506 | 2.67 | NA |
| GLN668 | GLN671 | 2.91 | NA |
| GLN671 | GLN668 | 2.92 | NA |

| A | B | Distance<br>model (Å) | Distance<br>exp (Å) |
| --- | --- | --- | --- |
| **ARG83** | **ASP183** | 2.61 | 8.74 |
| THR84 | SER210 | 2.81 | 5.95 |
| **LYS99** | **GLU247** | 2.59 | 7.74 |
| GLY106 | GLU209 | 2.81 | 6.01 |
| ASN140 | GLU247 | 2.71 | 4.92 |
| ASN140 | TYR248 | 2.81 | 6.26 |
| **ASP183** | **ARG83** | 2.59 | 7.38 |
| GLU209 | GLY106 | 2.9 | 8.09 |
| **GLU209** | **ARG238** | 2.61 | 3.14 |
| **ARG238** | **GLU209** | 2.6 | 3.1 |
| SER244 | LYS286 | 2.76 | 5.22 |
| **GLU245** | **LYS286** | 2.73 | 9.02 |
| GLU247 | ASN140 | 2.67 | 3.32 |
| TYR248 | ASN140 | 2.84 | 6.97 |
| LYS286 | SER244 | 2.85 | 6.39 |
| **LYS286** | **GLU245** | 2.77 | 7.1 |

**(2)Fzo1-(2)Ugo1** chain A (Fzo1) - chain B (Fzo1)
| A | B | Distance<br>model (Å) | Distance<br>exp (Å) | Prevalence<br>in simulation (%) |
| --- | --- | --- | --- | --- |
| ASN139 | ASP143 | 2.72 | 2.6 | 27.37 |
| LYS141 | LYS141 | 2.88 | 2.82 | 48.15 |
| **LYS141** | **ASP143** | 2.63 | 3.53 | 30.07 |
| ASP143 | ASN139 | 2.72 | 3.07 | 67.73 |
| ASP143 | LYS141 | 2.62 | 5.05 | 71.13 |
| GLN145 | SER171 | 2.89 | 3.02 | 85.71 |
| LYS149 | THR249 | 2.84 | 3.77 | 63.34 |
| LYS149 | GLU252 | 2.61 | 7.23 | 49.85 |
| ASN155 | LYS163 | 2.68 | 2.27 | 89.01 |
| **GLU157** | **LYS159** | 2.57 | 2.97 | 22.18 |
| **GLU157** | **LYS163** | 2.58 | 4.07 | 94.01 |
| **LYS159** | **GLU157** | 2.56 | 7.77 | 26.07 |
| LYS163 | ASN155 | 2.71 | 2.96 | 99.7 |
| **LYS163** | **GLU157** | 2.6 | 5.91 | 99.1 |
| SER171 | GLN145 | 2.91 | 3.24 | 9.49 |
| **ASP216** | **ARG380** | 2.57 | 3.08 | 2.2 |
| GLN217 | LYS374 | 2.94 | 2.91 | NA |
| GLN217 | ARG376 | 2.86 | 2.96 | 86.81 |
| **GLU252** | **LYS149** | 2.61 | NA | 3.5 |
| ASN318 | ASN318 | 2.95 | 2.97 | NA |
| ASN318 | LEU349 | 2.62 | 3.05 | 62.14 |
| ASN318 | SER350 | 2.75 | 2.88 | NA |
| MET319 | LYS352 | 2.69 | NA | 64.04 |
| **GLU344** | **LYS371** | 2.61 | 5.78 | 99.6 |
| LEU349 | ASN318 | 2.61 | 2.9 | 100.0 |
| SER350 | ASN318 | 2.81 | 3.29 | 2.6 |
| LYS352 | MET319 | 2.66 | NA | 85.41 |
| **LYS371** | **GLU344** | 2.6 | 5.21 | 95.3 |
| LYS374 | GLN217 | 2.88 | 3.37 | NA |
| ARG376 | GLN217 | 2.83 | 3.3 | 8.19 |
| **ARG376** | **ASP412** | 2.66 | 3.86 | 39.76 |
| **ARG380** | **ASP216** | 2.63 | 2.94 | 63.64 |
| **ASP523** | **ARG799** | 2.59 | NA | 2.1 |
| **LYS538** | **ASP777** | 2.72 | NA | 82.92 |
| **ASP777** | **LYS538** | 2.67 | NA | 36.76 |
| **ARG799** | **ASP523** | 2.6 | NA | 91.31 |
| **ARG803** | **GLU806** | 2.67 | NA | 26.47 |
| **GLU806** | **ARG803** | 2.67 | NA | 76.02 |
| **ASP810** | **LYS811** | 2.58 | NA | 2.6 |
| **LYS811** | **ASP810** | 2.6 | NA | 61.24 |

**Supplementary Table S4:**
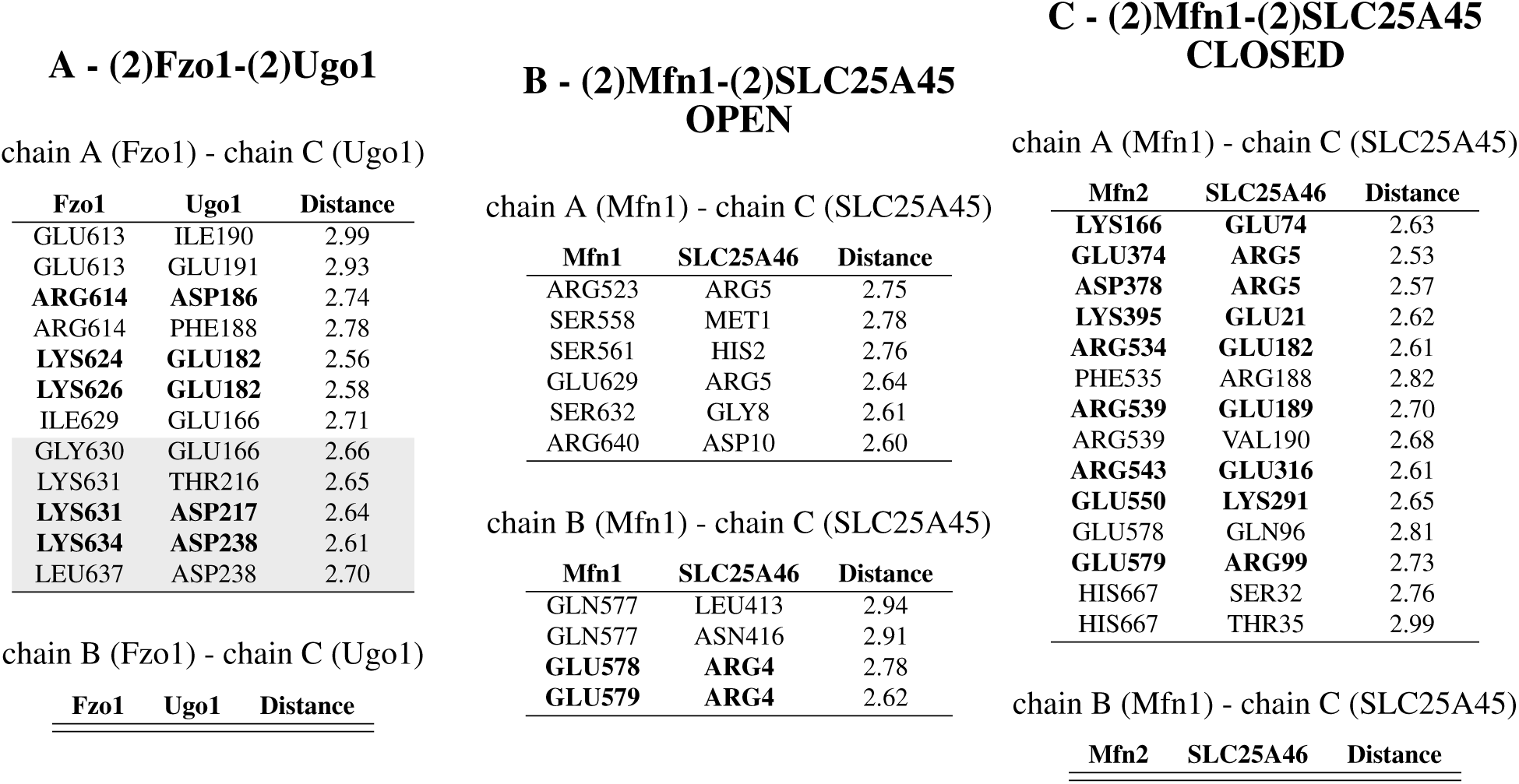
Contacts in Å between mitofusins and their solute carriers at a distance lower than 3Å. (A) Closest residues between Fzo1 and Ugo1 in the (2)Fzo1-(2)Ugo1 tetramer AF model. The grey lines indicate the regions of the proteins reported by Wong et al. [22] to be in contact with one another. (B) Residues at a distance lower than 3Å for Mfn1-(2)SLC25A46 of the Mfn1-Mfn2-(2)SLC25A46 tetramer AF model. (C) Residues at a distance lower than 3Å for Mfn2-(2)SLC25A46 of the Mfn1-Mfn2-(2)SLC25A46 tetramer AF model. Lines in bold represent the salt bridges.

**Supplementary Figure S8:**
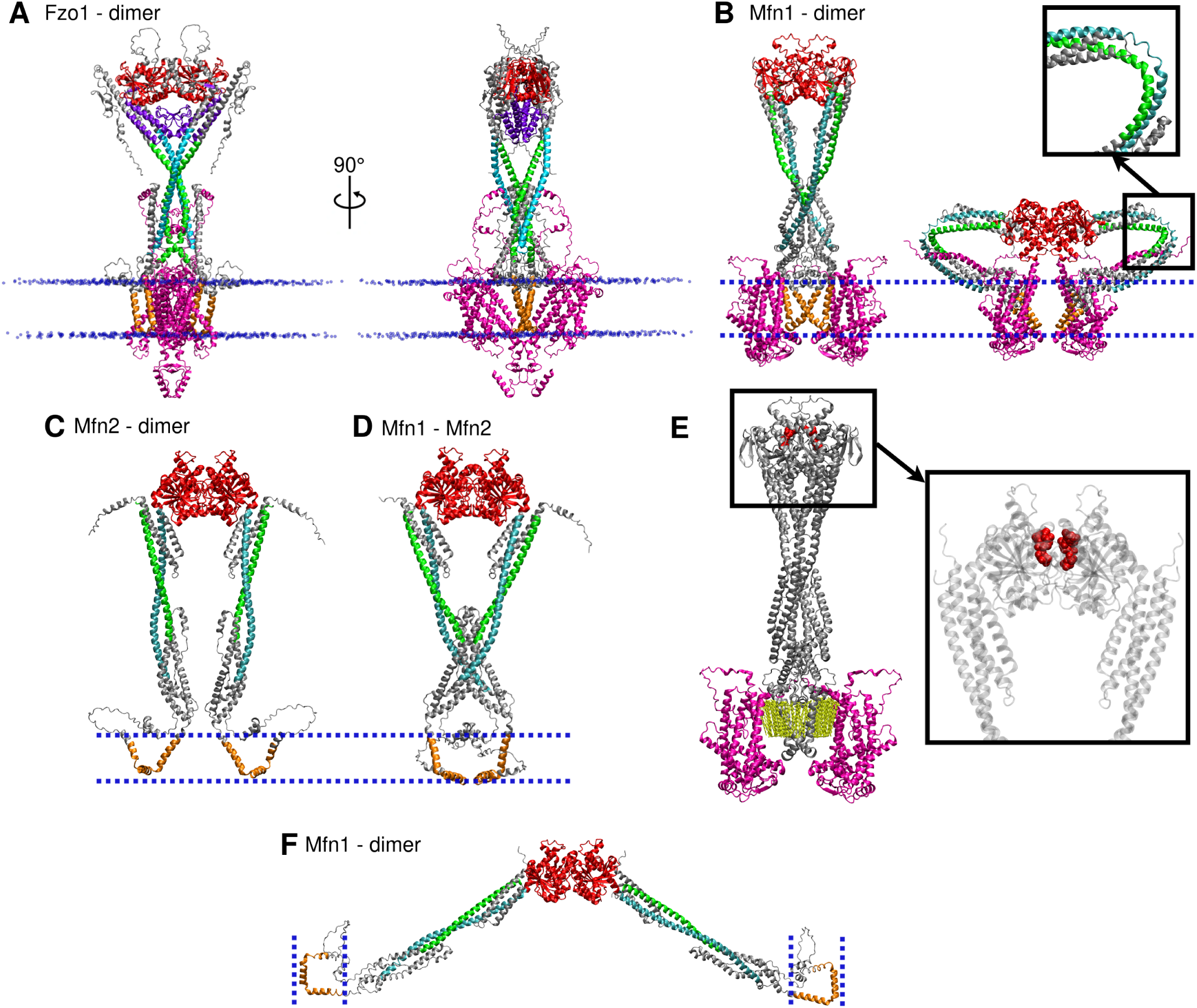
Complete AlphaFold models of mitofusins. (A) Heterotetramer of two Fzo1 and two Ugo1 produced by AF3 in the presence of oleic fatty acids and GTP, not shown here. The fatty acids molecules were systematically found located on the same plane as Ugo1. GTP was found binding to the GTPase domain. This model was used for the coarse-grained simulations. Fzo1 domains are colored as a function of the domains defined in 3(A). Ugo1 monomers are colored in magenta. (B) Heterotetramer of two Mfn1 and two SLC25A46 produced by AF3, in the presence of oleic fatty acids and GTP, not shown here. The fatty acids molecules were systematically found located on the same plane as Ugo1. GTP was found binding to the GTPase domain. Mfn1 domains are colored as a function of the domains defined in 3(A). SLC25A46 monomers are colored in magenta. (C),(D),(F) Respectively Mfn2 cis-dimer, Mfn1-Mfn2 cis-heterodimer and Mfn1 trans-dimer, colored as function of the domains defined in 3(A). (E) Heterotetramer of two Mfn1 (grey) and two SLC25A46 produced by AF3 (magenta), in the presence of oleic fatty acids (yellow licorice representation) and GTP (red Van der Walls representation).

**Supplementary Figure S9:**
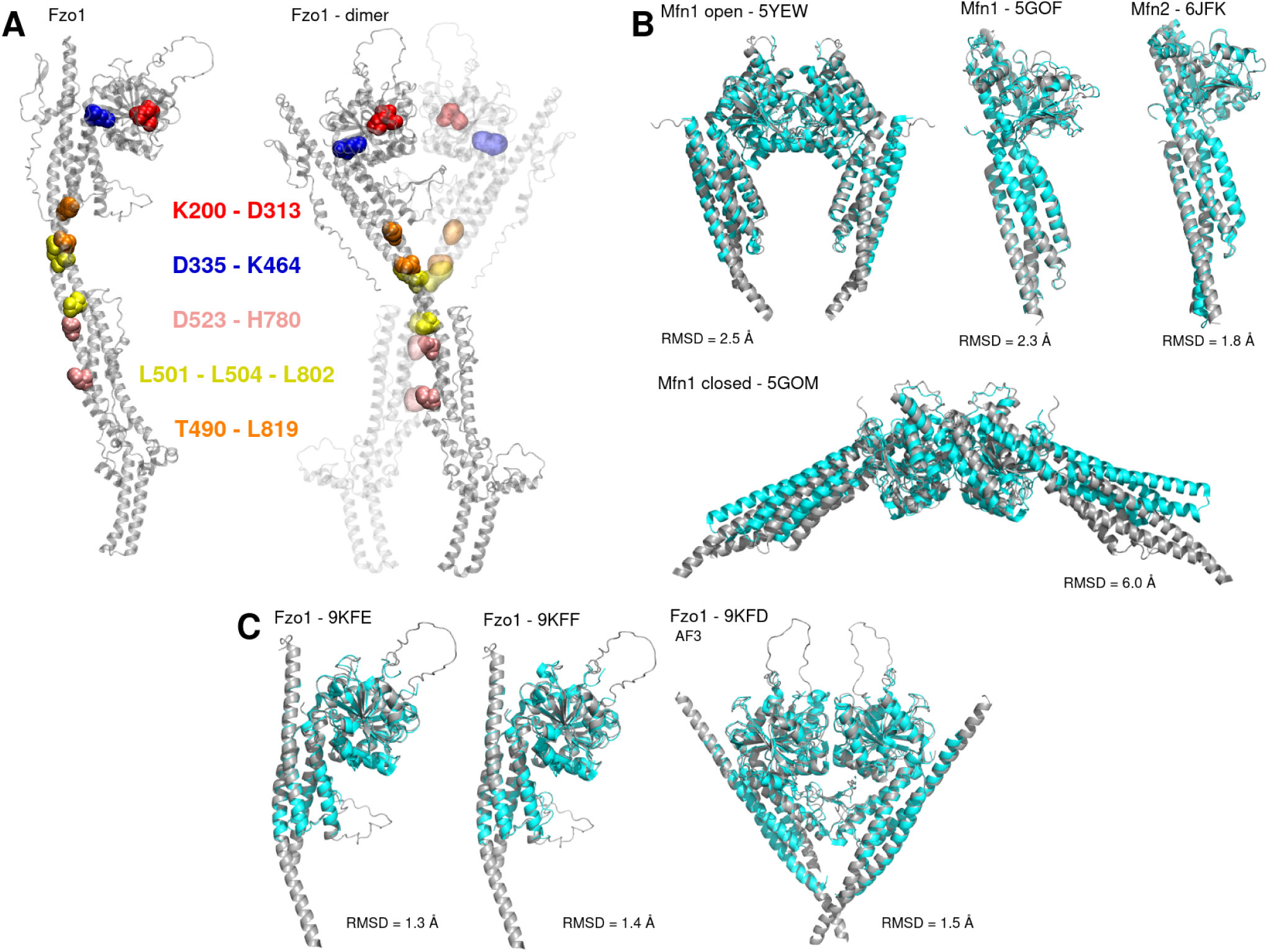
Experimental structures aligned with the Alphafold models. (A) Left, model obtained using AF. Right, model obtained using homology modeling [34, 35]. Residues in contact described in the 2017 model are shown in van der Walls representation and color-coded according to their respective groups of residues. (B),(C) In both panels the AlphaFold model are in grey, the experimental structure are in cyan. (B) AlphaFold prediction of Mfn1 and Mfn2 aligned with their solved structures. The open conformation (F3C) aligned with 5YEW has capri parameters corresponding to a medium quality model: fnat=0.67, irmsd=2.50 Å, dockQ=0.59. The closed conformation (F3D) aligned with 5GOM has capri parameters corresponding to a medium quality model: fnat=0.54, irmsd=2.67 Å, dockQ=0.35. (C) AlphaFold 3 prediction of Fzo1 aligned with its solved structures. The model in F3B aligned with 9KFD has capri parameters of : fnat=0.80, irmsd=1.50 Å, dockQ=0.75.

**Supplementary Figure S10:**
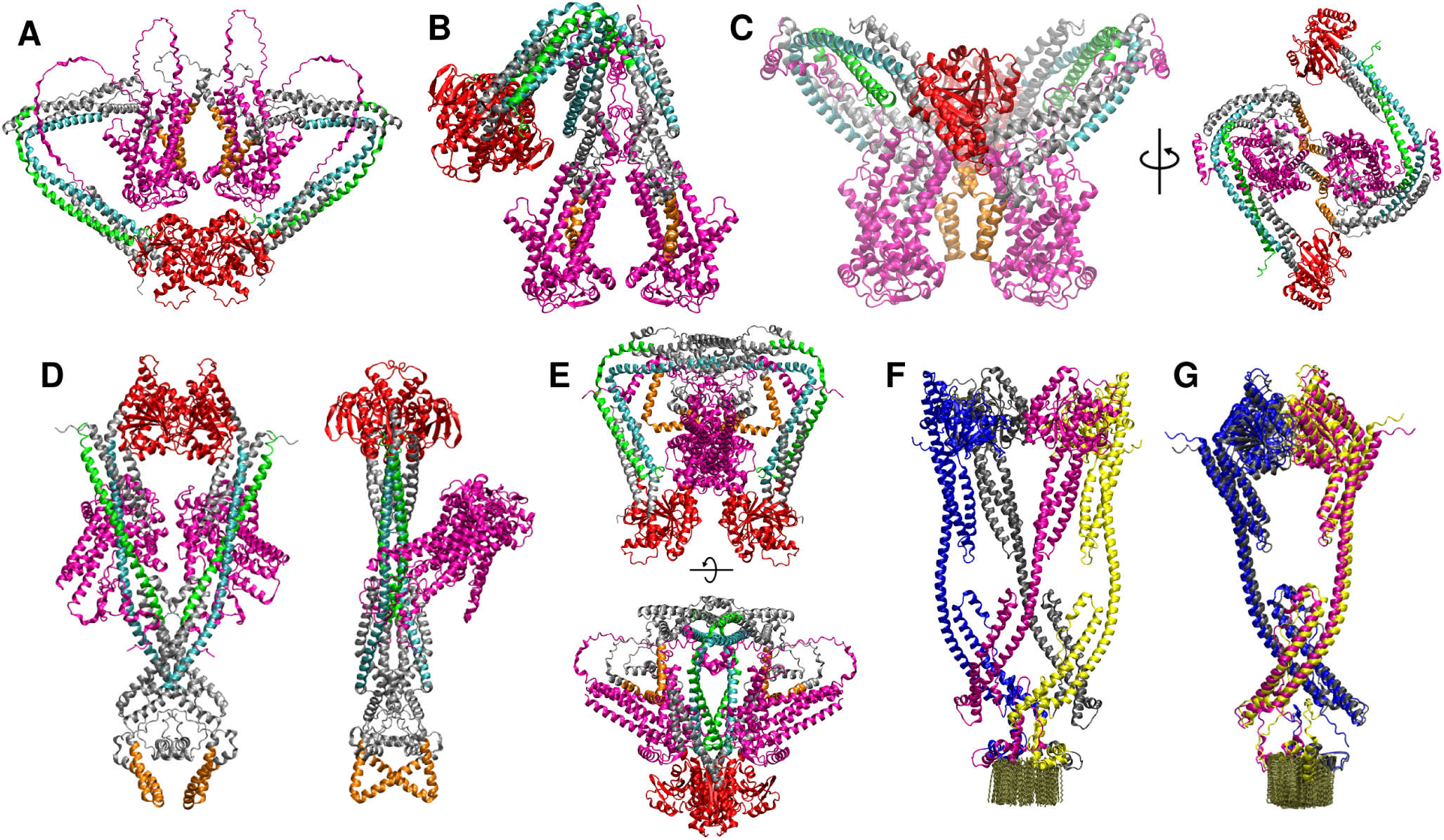
Diverse unrealistic conformations of Mfn1 produced by AlphaFold. (A),(B),(C),(D),(E) Heterotetramer of two Mfn1 and two SLC25A46 produced by AF3 in the presence of oleic fatty acids and GTP, not shown here. The fatty acids molecules were systematically found located on the same plane as SLC25A46. GTP was found binding to the GTPase domain. Mfn1 domains are colored as a function of the domains defined in 3(A). SLC25A46 is colored in magenta. (F),(G) Homotetramer of Mfn1 built with a mix of oleic and palmitic acids.

